# CF2H: a Cell-Free Two-Hybrid platform for rapid protein binder screening

**DOI:** 10.1101/2025.07.16.665152

**Authors:** Julien Capin, Pauline Mayonove, Angelique DeVisch, Amon Becher, Giang Ngo, Alexis Courbet, Robert J Ragotte, Martin Cohen Gonsaud, Julien Espeut, Jerome Bonnet

## Abstract

Protein binders that detect, activate, inhibit or otherwise modulate their targets are pivotal for biomedical applications. With the increasing accuracy and accessibility of *de novo* protein design, faster and cheaper experimental screening methods would democratize and accelerate the identification of high-affinity binders. Here we present Cell-Free Two-Hybrid (CF2H), a rapid and sensitive method for detecting high-affinity protein-protein interactions (PPI) that does not require cloning, protein purification nor high-end laboratory equipment. CF2H uses a dimerization-activated DNA binding domain (DBD) fused to prey and bait proteins to trigger transcription upon protein-protein interaction. We demonstrate that CF2H enables the detection of interactions between various types of target and binder proteins such as single-domain antibodies, DARPins and *de novo* designed binders. We benchmark CF2H as a screening platform by validating previously reported binders for Mdm2 and discover high-affinity binders targeting the checkpoint inhibitor PD-L1 in less than 24 hours. Finally, we show that CF2H can be used to characterize small-molecules modulators of PPI and detect protein biomarkers, opening the door for a new class of cell-free biosensors.

## Introduction

Protein-protein interactions (PPI) are involved in key biological processes such as signaling, cell fate, adhesion, and immunity^1,2^. Disruption of these interactions through genetic mutations or regulatory imbalances can lead to a wide range of diseases^3^. Targeting PPIs with therapeutic molecules, such as small compounds or engineered proteins, has proven effective in restoring molecular function, as exemplified by the clinical success of binders blocking immune regulatory checkpoints such as the cytotoxic T-lymphocyte-associated protein 4 (CTLA-4) or programmed cell death protein 1 (PD-1) pathways^4^. Specific protein binders such as antibodies and their single domain derivative are also invaluable tools for detecting, quantifying, and visualizing proteins in different cellular contexts. In synthetic biology, designing *de novo* protein binders with tailored interacting properties is crucial to engineer biosensors or develop orthogonal molecular circuits with predictable control over cellular behavior^5^.

Classical binder or antibody development has generally relied on resource and cost-intensive screening campaigns involving libraries containing billions of variants. This is beginning to change with AI-based protein design methods which aim to bypass these traditional high-throughput screening steps^6–8^. The design success rate of *de novo* protein binders has dramatically improved, reducing the number of candidates that must be experimentally tested from millions to just dozens or hundreds to obtain binders with reasonable affinities^6,9–11^. Moreover, significant efforts to open-source and democratize these tools have broadened the community of protein designers^10,12–14^. A major bottleneck now lies in the experimental validation of these designs— particularly for laboratories with limited biochemical expertise or insufficient access to adequate equipment.

So far, the methods of choice to validate *de novo* binders have been yeast display and automated protein purification pipelines followed by biophysical methods such as Surface Plasmon Resonance (SPR) or Bio-Layer Interferometry (BLI)^6,11^. Although yeast display is affordable and enables the screening of hundreds to thousands of binders, it requires initial burdensome steps for library construction and validation. High rates of false positives is a common drawback of this method, which can be reduced by increasing the number of sorting rounds involving extra culturing steps, or using a subtractive screening strategy, further increasing its experimental complexity. On the other hand, an automated purification pipeline followed by BLI or SPR is time-consuming and expensive, requires numerous washes and large amounts of purified proteins. Solution assays provide a more native environment, closer to the *in vivo* conditions, and require lower quantities of reagents.

We therefore sought to develop a one-pot solution assay for medium-throughput PPI screening that would be fast, cost-effective, and experimentally simple to remain accessible to non-specialized laboratories. We chose to harness the potential of Cell-Free Systems (CFS) to develop a Cell-free Two-hybrid assay (CF2H), enabling robust validation of *de novo* protein binders in hours rather than weeks.

A proof-of-concept PPI detection system was previously developed^15^ using the PURE^16^ system but this assay has a narrow operating range and cannot differentiate between interactions when the dissociation constant (K_d_) is lower than 10 nM, limiting its relevance for screening high-affinity binders. Moreover, despite recent progress towards home-made, cheaper alternatives^17^, the cost of PURE remains prohibitive for large-scale applications^18^.

We thus turned to *E.coli* lysate-based CFS, a cheap, convenient, and robust format for protein production^18–20^. Lysate-based CFS still contains the major chaperone systems found in bacteria and energy regeneration systems^21^, ensuring optimal production efficiency for a wide range of proteins^22,19,20^. We previously showed that using an *E. coli* strain deleted for RecBCD made the CFS compatible with protein expression from linear DNA templates, which is a critical aspect for developing a fast screening approach^23^.

We examined various options to develop a CFS-compatible PPI detection approach. We considered the use of split-protein complementation such as split-GFP, split-luciferase, or split-T7 polymerase but these systems present several limitations. All these assays are sensitive to fragment orientation, complicating fusion designs and necessitating the screening of various fusion types^24,25^. Split-luciferase assays rely on the addition of an exogenous substrate that terminates cell-free reactions, preventing access to real-time reaction dynamics. Split-GFP reassembly is irreversible, preventing access to dissociation constants (Koff), and potentially overrepresenting weak interactions^26^. Finally, the tendency of split-T7 fragments to spontaneously dimerize makes this system time-sensitive and prone to false positives^27^.

We opted for a two-hybrid approach whereby bait and prey proteins are fused to an inactive, monomeric DNA-binding domain (DBD). Upon bait and prey interaction, dimerization of the DBD restores its DNA-binding activity leading to the expression of a reporter gene. The cell-free assay that we developed is based on the CI transcriptional regulator from phage Lambda and its associated pRM promoter. We first show that CF2H can detect binding without non-specific background for a wide range of PPIs involving different protein binders such as VHHs, DARPins, and monobodies, all directly expressed from linear DNA templates. We benchmark our method on 48 de novo designed binders targeting human Mdm2 that were previously validated through a different screening approach^6^. We then demonstrate the translational value and accessibility of CF2H by discovering new high-affinity protein binders targeting murine PD-L1 and show their ability to block immune-checkpoint in a cellular model. Finally, we expand the applications of CF2H to test small molecule modulators of PPI and repurpose the system to perform small molecule and protein detection, demonstrating the potential of CF2H to engineer scalable cell-free biosensors. Together, our results establish CF2H as an accessible and flexible platform to rapidly screen *de novo* designed protein binders, assess small molecule modulators of PPIs and engineer clinically relevant biosensors.

## Results

### Development and optimization of a Cell-Free Two-Hybrid system

CI is a transcriptional regulator from the lambda bacteriophage that infects *Escherichia coli*, and controls the bistable switch between lysogeny and lytic growth by repressing the lytic promoters pL and pR while auto-activating its own promoter pRM to maintain lysogeny and superinfection immunity^28^. Early work on CI demonstrated its organization in two domains separated by a 40 AA linker, with the N-terminal domain (1-92) being involved in DNA binding while the C-terminal one (132-236) mediating dimerization and cooperative interactions between repressor molecules^29,30^ (**Fig1a**). CI N-terminus dimerization was then shown to be necessary for its DNA binding activity and transcriptional regulatory function^31^. Subsequent work showed that replacing the C-terminal domain of CI by homodimeric GCN4 leucine-zippers could restore its ability to dimerize, bind operator DNA and repress transcription in *E. coli*^32^. This chimeric system was then used to assess the impact of amino-acid substitutions by monitoring the immunity to phage superinfection conferred to *E. coli* by a functional repressor. Other studies used this bacterial integrated system to screen homo-and heterotypic PPI or dissociative inhibitors^33^, mainly using resistance to superinfection and repression of various reporter genes as a readout^34^. The simplicity of the CI interaction assay and its functionality in *E. coli* made it a candidate of choice for a lysate-based CFS PPI assay.

To engineer a circuit in which CI activates reporter gene expression, we constructed a reporter plasmid in which sfGFP is under the control of the pRM promoter. This promoter is activatable by CI through binding of both OR1 and OR2 operators. The third operator (OR3) was obliterated (OR3*) to prevent repression of pRM at high CI concentrations^35^. We then built a linear DNA expression cassette comprising a constitutive promoter, a RBS, and an engineered version of CI with an improved activation profile^35,36^, and added it to a cell-free reaction containing the reporter plasmid **(Fig1a)**. Once expressed, CI should homodimerize via its C-terminal domain, bind to operators OR1 and OR2, and stimulate transcription initiation through direct contact with the σ subunit of the RNA polymerase **(Fig1a)**. Robust expression of sfGFP was detectable within less than 2 hours at 37°C, with fluorescence intensity plateauing by 8 hours (**Fig1b**). This result confirmed that CI functions as a specific transcriptional activator of the pRM promoter when expressed from linear DNA fragment in lysate-based *E. coli* CFS. We then replaced the C-terminal domain of CI with the autodimerizing leucine zipper GCN4, as previously reported by Hu and colleagues^32^. We optimized the concentrations of the linear DNA encoding CI-GCN4 and the reporter plasmid, to an optimum producing a strong fluorescent signal, confirming that the system can function as a reporter of heterologous protein-protein interactions (**FigS1**).

**Figure 1:**
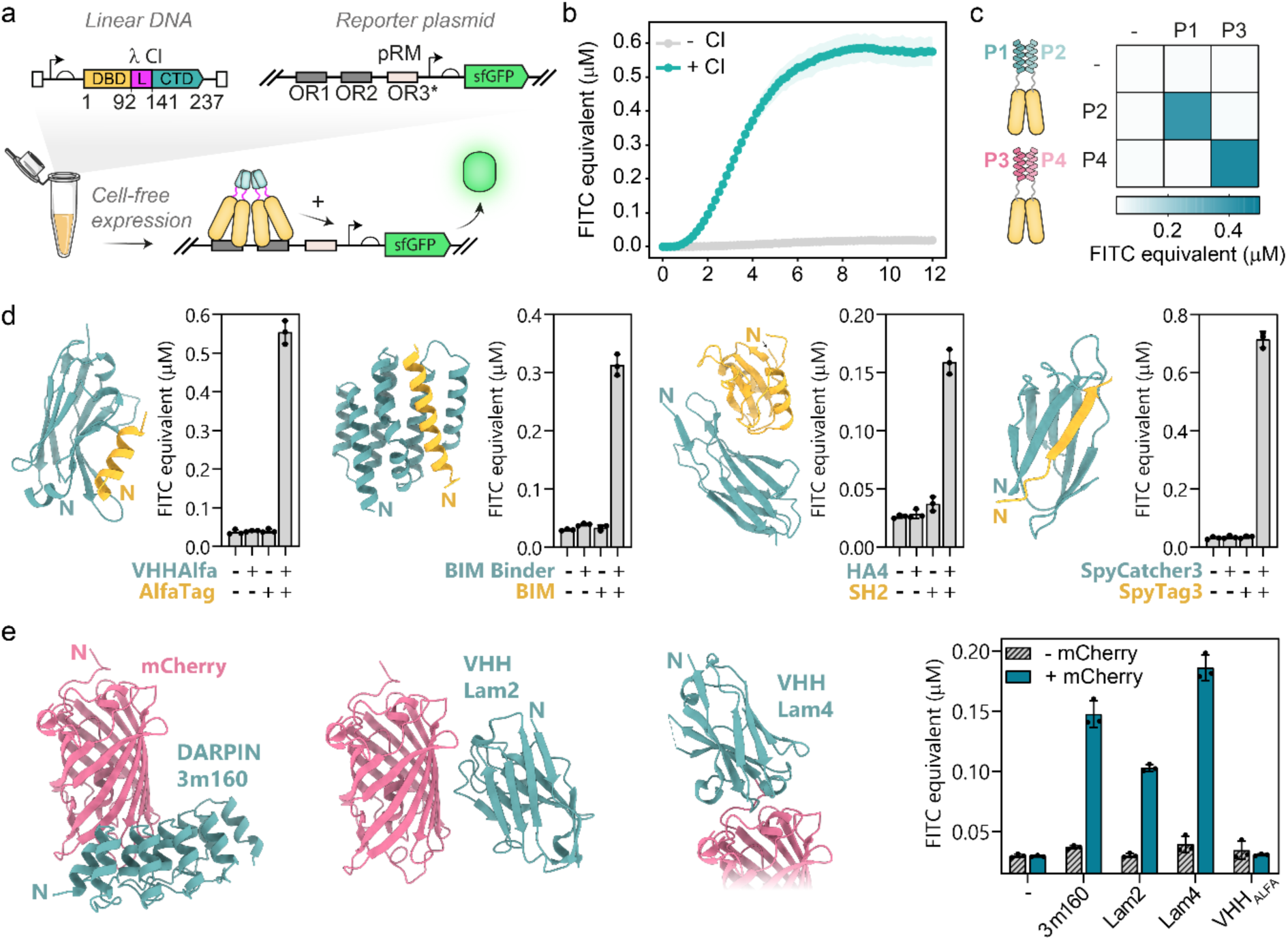
Engineering a Cell-free Two-Hybrid system based on Lambda repressor CI. **a.** Schematic representation of the Cell-Free two-hybrid system. A linear dsDNA fragment that contains spacer sequences (white boxes) used for PCR amplification, a constitutive promoter and RBS, as well as the lambda phage CI repressor coding sequence. CI comprises a DNA binding domain (DBD, gold), a linker region (L, magenta) and a C-terminal domain (CTD, light sea green) responsible for its dimerization. A reporter plasmid containing an sfGFP under the control of the CI-activated promoter pRM with a mutated OR3* operator to prevent self-inhibition at high concentrations of CI. **b.** Monitoring pRM activation by CI in CFS. Cell-free reactions containing the reporter plasmid were carried out in the absence or presence of CI-encoding linear DNA and monitored for 12 hours. Mean of 3 replicates +/-SD **c.** CI DBD (1-92) and the synthetic linker region (GGSx5) were fused to different coiled-coil pairs P1/P2 and P3/P4 to test the specificity of their interactions. Mean of 3 replicates. **d.** CF2H assays with various types of protein binders: VHH_ALFA_ (PDB 6I2G), de novo designed BIM binder (AF2 model), HA4 monobody (PDB 3K2M) or covalent binding with SpyCatcher3 (AF2 model). Mean at 8 hours of 3 replicates +/-SD. **e.** CF2H assays with DARPin 3m160 (AF2 model), VHH Lam2 (PDB 7SAJ), VHH Lam4 (PDB 7SAK) binding mCherry via distinct interfaces and different relative N-termini orientations. VHH_ALFA_ was used as a negative control. All bar graphs in d and e display the mean of 3 replicates at 8 hours +/-SD.

Next, we sought to optimize the CI DBD and linker regions. We first attempted to reduce the length of the DBD to its minimal form by removing 16 amino acids (CI 1-76) that were not directly involved in DNA binding. This shortened version of the DBD did not yield any activation regardless of the linker and PPIs tested (**FigS2, FigS3**). Direct fusion of proteins to the full-length DBD (CI 1-92) enabled signal production for some but not all protein couples tested. Adding a long flexible linker (GGSx5) greatly improved the response for larger globular protein domains such as mCherry or SpyCatcher (**FigS3**).

To assess the specificity of the assay, we challenged P1 and P2 to a second couple of orthogonal coiled-coils P3 and P4^37^. As expected, sfGFP expression was observed only in the presence of matching pairs, demonstrating that the signal observed in CF2H is driven by specific PPIs (**Fig1c**).

Next, we challenged our cell-free assay to detect various protein-protein interactions involving diverse synthetic protein binders (**Fig1d**). Of these, VHH antibodies are small (15 k_Da_) single antigen-binding domains of heavy-chain antibodies originating from camelids. VHH antibodies share a conserved disulfide bond, which requires periplasmic targeting or engineered *E. coli* strains for proper folding. Interestingly, lysate-based CFS produced from non-specialized *E. coli* strains show robust expression of soluble, functional VHHs^38^. VHH_ALFA_ binds a 15 amino acid helix named Alfa-Tag with picomolar affinity^39^. Co-expression of CI-VHH_ALFA_ and CI-Alfa-Tag led to a strong sfGFP expression in CF2H (**Fig1d**). Similarly, the binding of a *de novo* designed BIM binder to the BIM helical peptide could be detected in our system^40^. We next tested the HA4 monobody, another type of single domain antibody based on a synthetic scaffold from fibronectin III domain (FN3)^41^. This sdAb binds the SH2 domain of human cellular tyrosine protein kinase Abl1 with nanomolar affinity. Although the level of signal observed was lower than the couples tested previously, this interaction was clearly detectable in CF2H (**Fig1d**). Finally, we included a specific type of interaction involving a *Streptococcus pyogenes* originating adhesin protein, split into a Catcher and a Tag and further engineered to form an intermolecular covalent bond upon binding^42^. Indeed, this irreversible interaction led to the highest sfGFP expression overall (**Fig1d**).

Finally, we investigated whether the relative spatial orientation of target and binder proteins could potentially hinder their detection in CF2H. For PPI assays relying on direct protein fusions, steric clashes can come into play and give rise to false negatives. Using mCherry as a target protein, we tested 3 binders with distinct interaction surfaces : DARPin 3m160^43^, VHH Lam2 and VHH Lam4^44^ (**Fig1e**). All three binders led to a substantial production of sfGFP expression only in presence of mCherry while the VHH_ALFA_ control did not, demonstrating that the flexibility in the linker region can tolerate interactions with a range of target/binder spatial orientations.

All together, these results demonstrate that protein heterodimerization can be monitored in lysate-based CFS via fusion with CI DBD, and that CF2H performs robustly across a broad range of target and binder proteins.

### Experimental validation of *de novo* protein binders

Next, we sought to apply CF2H to screen computationally designed protein binders. To benchmark our approach on a realistic number of candidate proteins, we tested 48 designs produced by Watson and colleagues and measured their interaction with human Mdm2^6^. Of these 48 candidate proteins, we selected 36 designed to target Mdm2, based on their diverse binding scores obtained by single-point BLI (SP BLI). Since the reported success rate on this particular target was exceptionally high (>50%), we also included 12 binders that were originally designed to bind an unrelated protein, IL-7Ra, as negative controls^6^. Upon reception, synthesised DNA fragments were amplified by PCR and used directly in CF2H (**Fig2a**).

**Figure 2.**
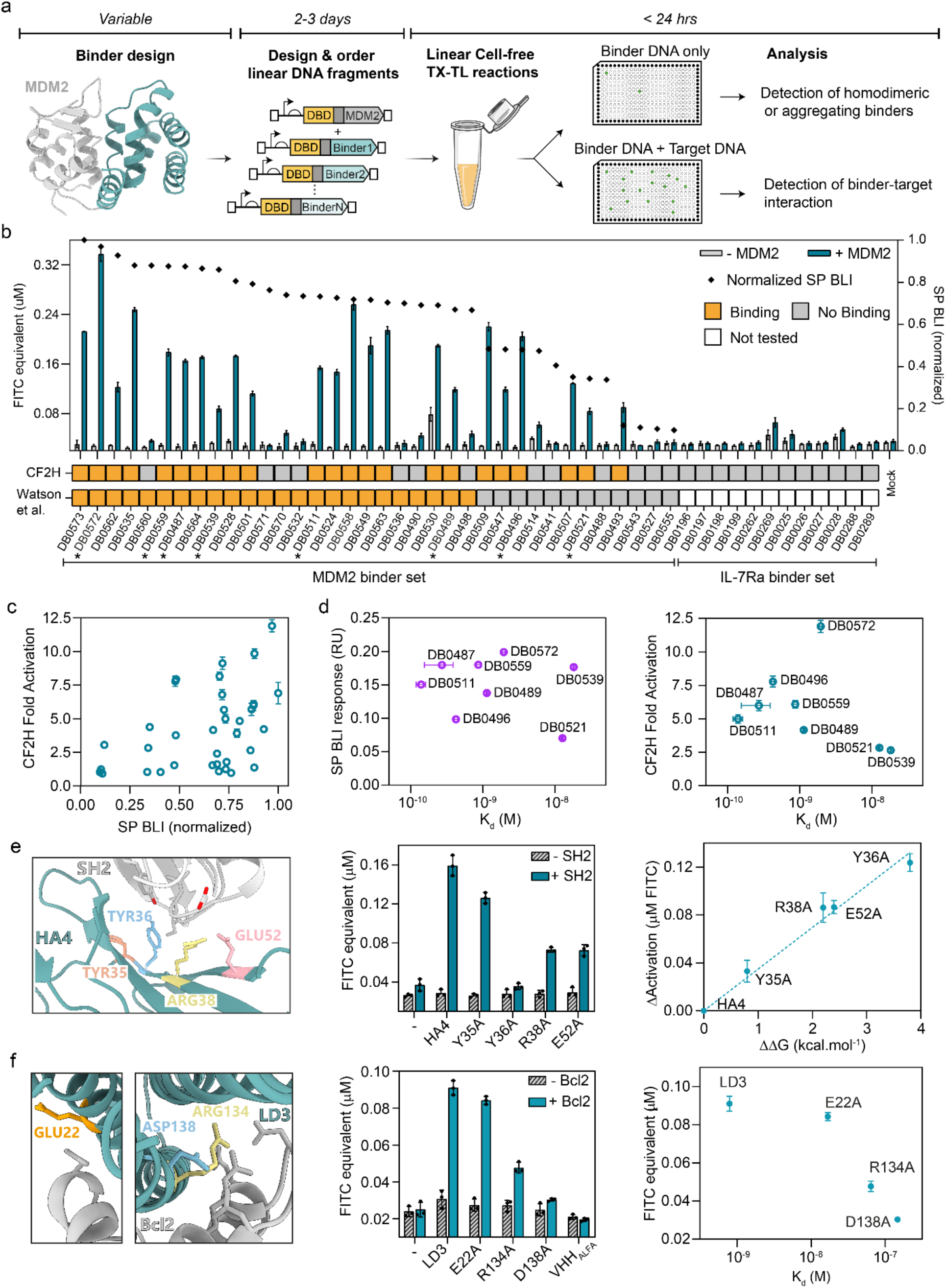
Benchmarking the CF2H workflow for screening *de novo* designed protein binders. **a.** Workflow of CF2H applied to *de novo* binder validation. All DNA constructs were purchased as linear double-stranded fragments. Upon reception DNA fragments were PCR amplified, concentrations were normalised and CF2H assays were carried out and analysed within 24 hours. **b.** Screening of Mdm2 *de novo* protein binders from Watson et al.^6^. 36 designs from the Mdm2 binders set and 12 binders from the IL-7Ra set were tested in absence or presence of Mdm2. A candidate protein was classified as a binder if it exhibited more than a twofold increase in signal upon incubation with its target compared to the background signal in the absence of target. Normalized single point BLI measurements (◆) were used to rank binders from left to right by decreasing affinity. Binders for which a K_d_ value was determined are shown with a (*) **c.** Correlation between single point BLI scores and fold activation in CF2H. **d.** Correlation between signal intensities and binding affinities for SP BLI and CF2H fold activation on eight Mdm2 binders for which K_d_ values were determined. **e.** Alanine scanning analysis of the HA4 interface residues interacting with the Abl1 SH2 domain using CF2H agrees with previously reported ΔΔG values (right panel). **f.** Bcl2-targeting *de novo* designed binder LD3 (PDB 6IWB) and alanine mutants with lower affinities were tested. Their activation profiles were compared to previously determined affinity values (right panel). All panels display the mean of 3 replicates +/- SD.

Since protein–protein interactions can exhibit a wide range of affinities, it is challenging to define a protein binder in binary terms. However, when screening, it is often convenient to set a threshold to focus downstream characterization on the most promising candidates. Watson and colleagues arbitrarily defined this threshold at 0.5 of normalized BLI score (**Fig2b**), meaning all the proteins above this value were binders. In CF2H, we classified a protein as a binder if it exhibited more than a twofold increase in signal upon incubation with its target compared to the background signal in the absence of target. Out of 24 proteins that were classified as binders by SP BLI^6^, 17 could be confirmed as binders in CF2H (**Fig2b**). Interestingly, DB0530 led to significant sfGFP expression even in absence of Mdm2, which may indicate its tendency to dimerize or aggregate. Moreover, of the 12 proteins defined as non-binding (SP BLI < 0.5), we found that 50% were binders in CF2H. Finally, none of the IL-7Ra binding proteins displayed any significant signal in presence of Mdm2, confirming the specificity of the screen. Direct comparison between CF2H fold-activation and SP BLI response values obtained by Watson and colleagues showed similar trends (**Fig. 2c**).

These experiments demonstrate that the CF2H workflow enables reliable screening of *de novo* protein binders. In less than 24 hours, the assay provides comparable information as conventional characterization assays that require weeks, expertise and specialized equipment.

### Influence of binding affinity on CF2H activation levels

Amongst the Mdm2 binders tested, dissociation constants were determined for eight of them. We compared these affinity values to the response intensities obtained by single-point BLI or CF2H (**Fig2d**). We observed that single-point BLI does not provide any additional information on the strength of the interaction. In CF2H, it is expected that in addition to affinity, other factors such as differences in expression levels, stability, and solubility may contribute to the varying signal intensities. Nonetheless, we could distinguish weaker binders that are in the tens of nanomolar range (DB0521 and DB0539) from higher affinity binders with nanomolar or subnanomolar K_d_. To explore whether CF2H is indeed sensitive to differences in binding affinity, we decided to perform alanine mutagenesis assays. In this case, a single amino acid is substituted at a time, which should limit the impact of differential expression, solubility and stability on the signal intensity. We first exposed the SH2 domain of Abl1 to 4 different alanine mutants of the HA4 monobody previously described^41^. While mutating tyrosine 35 had a limited impact on binding, mutating the adjacent tyrosine completely abolished the response (**Fig2e, FigS4**). Substituting the two other residues R38 and E52 decreased the level of fluorescence produced by around half. Interestingly, the difference in signal measured between these mutants and HA4 (ΔActivation) in CF2H correlates with ΔΔG values previously obtained by Surface Plasmon Resonance (SPR), confirming that CF2H can provide some insight on the impact of individual mutations for a given interaction. In the same vein, we measured the interactions between human Bcl2 and alanine mutants of the computationally designed LD3 protein^45^. We observed that the E22A mutation barely affected the response compared to LD3, while R134A and D138A had the highest impact on the binding in CF2H (**Fig2f, FigS4**). When comparing these values to the previously determined affinities we observed that CF2H, despite not accurately recapitulating the impact of E22A on Bcl2 binding, could still rank the mutants according to their binding affinities (**Fig2f**).

These results indicate that CF2H is sensitive to variations in binding affinity, a relevant feature for the identification of high-affinity binders.

### Discovery and characterization of checkpoint inhibitors targeting PD-L1

We next sought to screen a set of in-house–designed binders targeting PD-L1, a key molecule in immune checkpoint blockade^4^, for which *de novo* binders could help the development of new cancer immunotherapies. These binders were generated against the extracellular domain of the murine homolog of PD-L1 (mPD-L1) using BindCraft — a recently introduced open-source pipeline that leverages AlphaFold2 backpropagation for one-shot binder design^10^. The streamlined BindCraft pipeline aligns well with the philosophy of our screening workflow, offering user-friendly features that make it accessible to researchers without specialized expertise in protein design.

We also chose PD-L1 because it is a challenging target - its overexpression in *E. coli* results in misfolding and the formation of inclusion bodies^46^, and expression in lysate-based CFS had, to our knowledge, not been reported. Using CF2H, we first attempted to validate the interaction between mPD-L1 and previously described VHH antibodies. However, we observed that expression of CI-mPD-L1 alone led to high levels of activation of the pRM promoter, likely due to aggregation of the mPD-L1 domain (**FigS5a**). We then wondered whether we could modify the CF2H system to enable PPI detection despite part of the target protein forming aggregates.

To avoid pRM activation caused by target aggregation, we moved away from the CI-mPD-L1 fusion. Instead, we sought to link CI-DBD only to the binder while promoting target multimerization by fusing mPD-L1 to a homo-multimeric domain. Thus, upon recognition of multimeric mPD-L1, CI-linked binders should multimerize and activate transcription. We hypothesized that this modified approach would decrease background signal and preserve functionality through target-induced multimerization. We sought to test different multimerization domains: homodimeric GCN4 leucine zipper^47^, parallel tetrameric coiled coils (CC-tet)^48^, tetramerization domain of p53 (Tp53)^49^, hexameric parallel coiled-coils (CC-Hex)^50^ and an heptameric mutant of GCN4 (GCN4-hept)^51^ by fusing them to the N-terminal of the target protein, keeping the GGSx5 linker for flexibility. We first tested these domains on Mdm2, a target that we previously characterized, in order to compare the performance of multimer-Mdm2 to CI-Mdm2 systems. Interestingly, we observed that with the exception of GCN4, all multimerization systems produced at least as much signal as the CI-Mdm2 approach (**FigS5b**). Amongst these, Tp53 and CCHex led to the highest signal, around 3-fold higher than CI-Mdm2. We subsequently applied this multimerization approach (herefrom referred to as CF2H-multimer) to mPD-L1 using the Tp53 domain and could detect interaction with all the VHH tested (**FigS5b**).

We then used CF2H-multimer to screen 36 newly designed mPD-L1 binders selected amongst a hundred designs generated using the BindCraft pipeline (**FigS6**)^10^. Although only low levels of GFP were measured, binders DBP001 and DBP035 displayed substantial activation only in presence of multimeric mPD-L1 (**FigS7**). We also observed high levels of homodimerization or aggregation for two of the binders, DBP026 and DBP043, two candidates predominantly composed of beta strands (**FigS6**).

While these results were promising, we sought to enhance the signal-to-noise ratio of the PD-L1 screening assay. We reasoned that spiking in some purified mPD-L1 in CF2H reactions instead of producing it in CFS could help enhance the signal. Similarly to the CF2H-multimer approach, the purified mPD-L1 protein had to be in a multimeric form for the system to work. Hence, we used a biotinylated version of mPD-L1, which we incubated with streptavidin prior addition into CF2H reactions. Streptavidin naturally assembles into a tetramer, with each subunit capable of binding one biotin molecule^52^. We reckoned that coupling biotinylated mPD-L1 to streptavidin would result in mPD-L1 tetramers that would be conducive to CI-multimerization and transcriptional activation in case of CI-linked binder interaction (**Fig3a**).

We validated this strategy probing the interaction between anti-mPD-L1 VHH antibodies and pre-assembled Strep-mPD-L1 complexes spiked into CF2H reactions. We observed a dose-dependent signal for all VHHs only upon addition of streptavidin-complexed mPD-L1 (**Fig 3a, FigS8**). We confirmed the scalability of this approach by testing VHHs targeting the immune checkpoint proteins CTLA-4 (PDB 5E03) and CD47^54^ with sub-nanomolar affinity (**FigS9, FigS10**).

Encouraged by these results, we harnessed this approach to repeat the screen of the mPD-L1 candidate binders. We observed a significant increase in GFP signal for 7 out of the 36 binders tested (**Fig3b**). Interestingly, DBP001 and DB035 produced the highest levels of activation in this screen, in line with the results obtained with the Tp53-mPD-L1 approach (**FigS7**). We also observed consistent levels of expression for most of the binders tested (**FigS11a**). Next, we confirmed binding of DBP023 and DBP035 by SPR, highlighting high-affinity binders, with Kd of 6.85 nM, and 0.78 nM, respectively (**Fig3c**). Although the affinity of DBP001 was confirmed by ELISA and SPR, the quality of the fitting was not as convincing as for the other binders (**FigS11c,d**).

**Figure 3.**
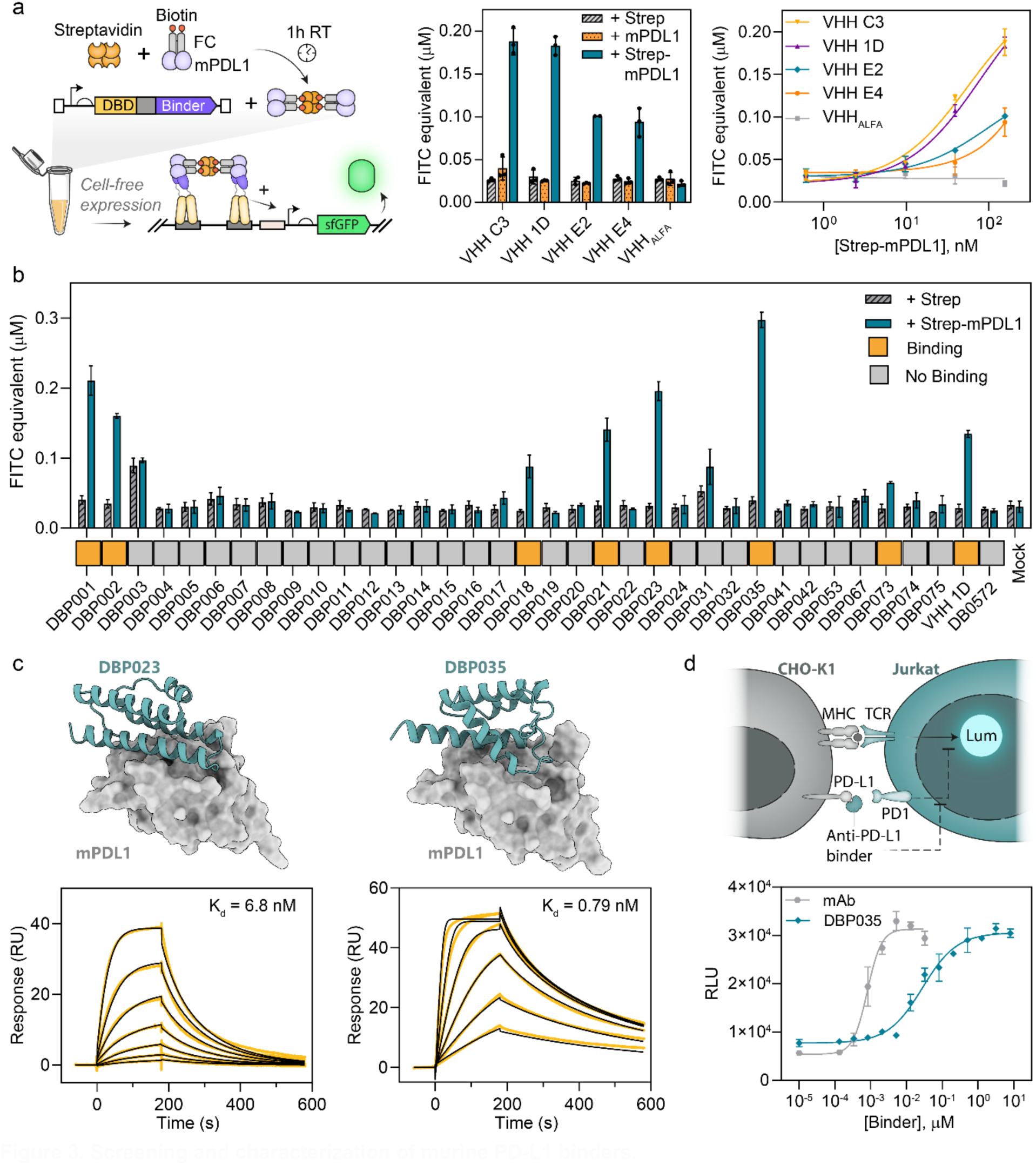
Screening and characterization of murine PD-L1 binders. **a.** Schematic representation of the multimerization of biotinylated mPD-L1 mediated by incubation with streptavidin before addition into CF2H. Streptavidin, mPD-L1 or streptavidin-mP-DL1 (1:1) was added to CF2H reactions expressing CI-fused mPD-L1-targeting VHHs and VHH_ALFA_ (middle panel). Dose-response behaviour of the CI-bound VHHs subjected to varying concentrations of strep-mPD-L1 (right panel). VHH-1D from PDB: 5DXW, all others from Broos et al.^53^). FITC equ. at 8 hours. Mean +-SD of 3 replicates. **b.** This strategy was exploited for rapid screening of mPD-L1 binders. Data shows FITC equ. at 8 hours. Mean +-SD of 3 replicates. For clarity, results for DBP026 and DBP023 are presented in FigS11b. **c.** AF2 models of DBP023 and DBP035 in complex with mPD-L1 and binding affinities determined by SPR. Response to injections of binders at 0.312 nM to 20 nM for DBP023 and 0.625 nM to 20 nM for DBP035 are shown in gold and corresponding fit in black. **d.** Determination of immune checkpoint blockade activity of DBP035. CHO-K1 cells expressing mPD-L1 were co-cultured with Jurkat cells displaying mPD-1 and harbouring a luciferase gene regulated by the Nuclear factor of activated T-cells (NFAT) pathway. Cells were exposed to increasing concentrations of DBP035 or anti mPD-L1 monoclonal antibody (mAb), and luminescence was measured after 6 hours of incubation. Mean +-SD of 3 replicates.

Next, we investigated the functionality of DBP035 by testing its ability to block the PD-1/PD-L1 immune checkpoint in a cellular model. Chinese Hamster Ovary cells (CHO-K1) overexpressing mPD-L1 were co-cultured with Jurkat cells displaying mPD-1. In this model, Jurkat cells harbor a luciferase reporter cassette linked to the Nuclear factor of activated T-cells (NFAT) pathway, which is suppressed by PD-1/PD-L1 interaction. Disruption of PD-1/PD-L1 interaction results in increased levels of luminescence. DBP035 addition to the cell culture was able to disrupt PD-1/PD-L1 interaction in a dose dependent manner, with a half-maximal inhibitory concentration (IC50) of 28 nM (**Fig3d**).

These results demonstrate that CF2H is a robust and modular platform for screening *de novo*-designed binders. For target proteins that are difficult to produce in CFS, purified streptavidin-templated antigens can be used as targets triggering multimerization of CI-linked binders. Finally, the CF2H pipeline enables the generation of functional binders, as illustrated by the ability of DBP035 to disrupt the PD-1/PD-L1 immune checkpoint interaction, emphasizing the translational value of the platform.

### Detection of PPI modulations

Protein-protein interactions can be modulated by the addition of small molecules, which can either break or stabilize interactions through binding in specific pockets. Such molecules represent more than 60% of FDA-approved drugs yearly^55^. Hence, PPI assays that can be extended to serve as screening platforms for competitive inhibitors, molecular glues or other small molecule modulators of PPIs are highly valuable.

We sought to challenge CF2H to monitor the effect of various types of small molecule modulators of PPIs. First, we tested the Mdm2 inhibitor NVP-CGM097, which binds Mdm2 and blocks the interaction with the p53 helix^56^. Since the *de novo* binders were scaffolded around the p53 helix, we reasoned that this drug should also perturb their interaction with Mdm2. We selected 3 different binders that displayed varying levels of activation DB0572, DB0558 and DB0489 and included the VHH_ALFA_ / Alfa-Tag couple as a control of specificity. Interestingly, we observed that only the binder DB0489, which displayed the weakest signal in CF2H (**Fig2b**), could be outcompeted at the two highest drug concentrations (**Fig4b**), validating experimentally its interaction interface with Mdm2. In the conditions tested, the other two binders were not affected by the drug.

Similarly, we subjected BCL2/LD3 to increasing concentrations of Venetoclax, a Bcl2 inhibitor^57^. We observed that the BCL2/LD3 interaction could be inhibited by Venetoclax with an IC50 ∼ 202 nM and that this inhibition was specific as the binding between VHH_ALFA_ and Alfa-Tag was not affected (**Fig4c**).

**Figure 4.**
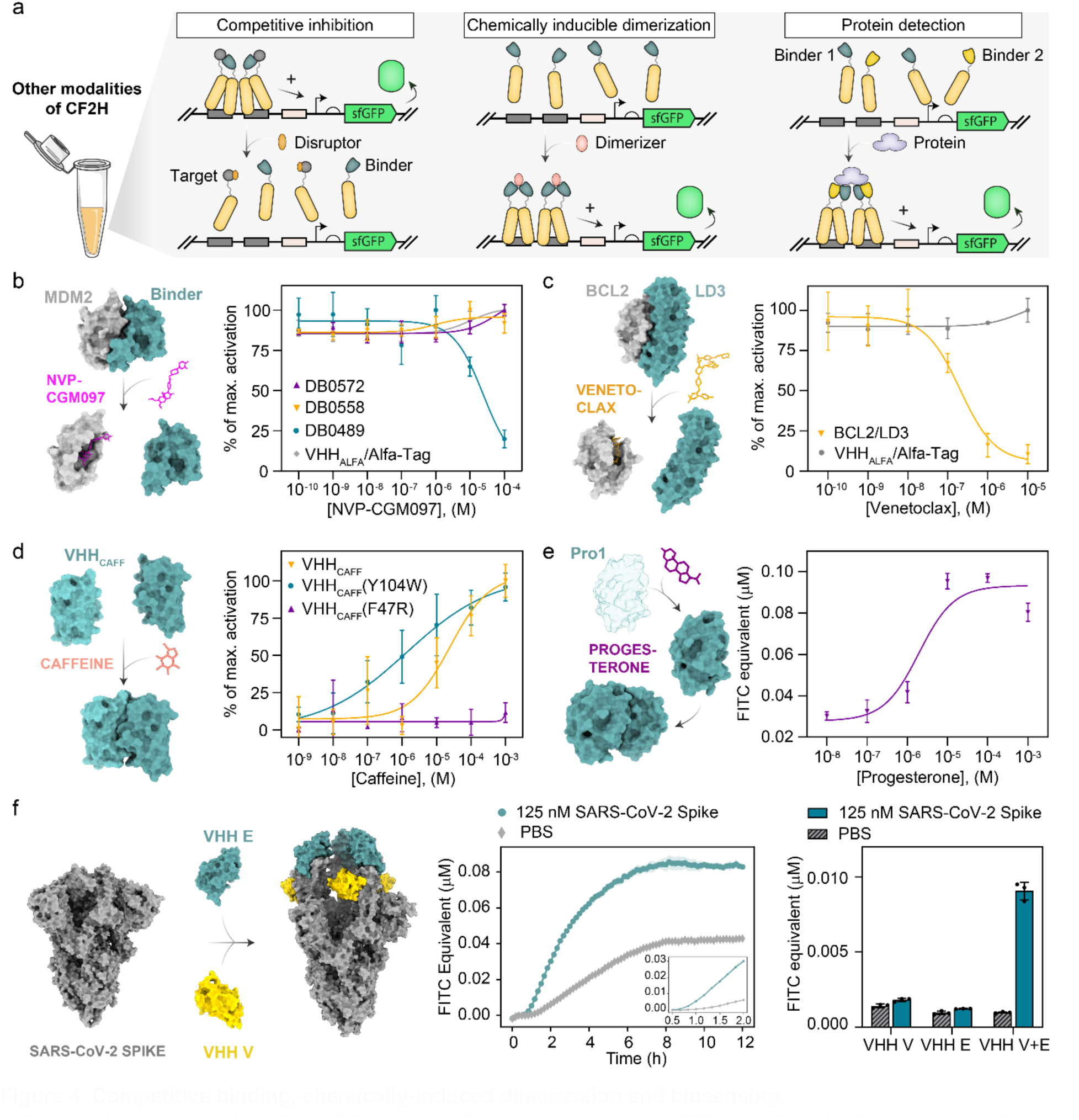
Competitive binding, chemically-induced dimerization and biosensing. **a.** Schematic representation of the different modalities possible with the CF2H platform. **b.** Competitive inhibition of Mdm2/de novo binder interaction with NVP-CGM097, an inhibitor of Mdm2. Increasing concentrations were added to CF2H reactions expressing Mdm2 and p53-inspired de novo binders. VHH_ALFA_/Alfa-Tag interaction was used as a control. **c.** Similarly, the BCL2-LD3 interaction was challenged by increasing concentrations of Venetoclax, a potent BCL2 inhibitor (PDB 6O0K). VHH_ALFA_/Alfa-Tag interaction was used as a control. **d.** Chemically induced dimerization of a single domain antibody (VHH_CAFF_) upon addition of caffeine (PDB 6QTL). Mutants with reported increased or abolished sensitivity to caffeine Y104W and F47 respectively, could be confirmed in CF2H. **e.** An evolved progesterone binding protein (Pro1) is stabilized by progesterone, enabling its homodimerization (Chai-1 model^69^). Mean of 3 replicates +/- SD. **f.** Detection of SARS-CoV-2 Spike protein trimer by simultaneous expression of CI-VHHV and CI-VHHE targeting neighbouring Receptor Binding Domain (RBD) epitopes (middle panel) compared to VHH V and VHH E expressed alone after 1 hour (Right panel). Mean of 3 replicates +/- SD. sfGFP production kinetics corresponding to these results are presented in FigS12.

Together, these results demonstrate that CF2H offers a fast and reliable method for screening competitive small-molecules modulators of PPIs.

### Biosensor engineering

Although the majority of small molecules developed for therapeutic applications are inhibitors of PPIs, other chemicals induce or stabilize dimer formation. Chemically inducible dimerization (CID) systems are broadly used in cellular biology as they offer a way to timely control the association between proteins of interest^58,59^. Ligand-induced dimerization leading to reporter activation is also a practical and robust method for biosensor engineering, *in vitro* and *in vivo*^27,45,59–63^. We thus sought to explore the potential of CF2H as a novel platform for biosensing applications. We first aimed at demonstrating small-molecule biosensing by using as a sensing domain a VHH that forms homodimers in the presence of caffeine^64,65^. We previously leveraged this system to engineer a caffeine-inducible switch in *E. coli*^61^. Other studies have shown that VHH_CAFF_ single mutants Y104W and F47R resulted in an increased or abolished sensitivity to caffeine, respectively^66,67^. We decided to test all three proteins in CF2H and showed that we could not only detect the effect of caffeine on dimerization of VHH_CAFF_ but also the impact of mutations on the system’s sensitivity **(Fig4d**).

Beyond direct PPI modulation, small molecule binding can also affect protein stability. This is the case for the progesterone-binding domain (Pro1) initially designed for ligand-dependent transcriptional control in yeast^68^. In the absence of progesterone, this protein domain cannot fold into a stable structure, which prevents its homodimerization. Progesterone acts as a pharmacological chaperone and stabilizes Pro1. In yeast, 1 µM of progesterone was necessary to see detectable transcriptional activation while 10 µM gave the maximum response. Interestingly, we obtained comparable results in CF2H, with an EC50 of ∼1.9 µM (**Fig4e**).

Finally, after validating small-molecule detection, we challenged CF2H to detect proteins, which are highly important diagnostics targets^70^. We chose the spike protein of the Severe Acute Respiratory Syndrome Coronavirus 2 (SARS-CoV-2)^71^ as a clinically relevant target. VHH_E_ and VHH_V_ are single domain antibodies targeting neighbouring epitopes of the spike receptor binding domain (RBD)^72^, which were previously used in a split-biosensor assay^27^. By combining the expression of CI-fused VHH_E_ and VHH_V_, we observed a rapid and specific increase in GFP signal in presence of Spike protein, with a clear unambiguous spike detection was possible after 1 hour, demonstrating the functionality of our biosensor (**Fig4f**). Although the activation level was not high, the difference in signal compared to the control was maintained across 12 hours. Interestingly, expressing VHH_E_ or VHH_V_ separately was not sufficient for sensing the spike trimer. These results are in line with results obtained in **FigS5b,** in which GCN4-Mdm2 failed to activate the system, suggesting that the formation of higher-order CI multimers may be more favourable for protein biosensing applications.

These data show that CF2H can respond to chemically-inducible dimerization and supports the engineering of cell-free biosensors for small-molecule and protein targets.

## Discussion

Here we developed CF2H, a rapid and simple workflow for high-affinity binder screening. Our system was designed with the aim of making binder screening simpler, more affordable and accessible to the community. By combining an *E. coli* lysate-based cell-free system compatible with expression from linear DNA templates to a two-hybrid approach, we constructed a workflow that bypasses tedious cloning, culturing and sequencing steps. Our experimental setup only requires a set of pipettes and a microplate reader for GFP measurements, and its complexity is comparable to setting up PCR reactions.

We showed that CF2H could provide comparable information as some of the currently employed screening methods relying on heavy automated purification pipelines followed by single point BLI measurements. One of the strengths of CF2H is the rapidity of the assay, with data obtained in hours rather than weeks. This makes this system particularly well-suited for rapid iterative design-test cycles, which could be used for affinity maturation or for developing active learning models.

Our *E. coli* lysate-based assay offers a cost-efficient platform, with an estimated expense of €0.10 per reaction^18^ (excluding DNA synthesis), bringing the overall cost of a standard de novo binder screen to under €10 using a home-made extract. For laboratories that would rather purchase CFS, the price per reaction ranges from €5-10 depending on the manufacturer. This cost remains highly competitive compared to other screening methods such as yeast display or automated purification and characterization workflows, especially when taking labour into account. Similarly to other Transcription-Translation (TXTL) platforms, CF2H automation would be straightforward and could be combined with acoustic liquid handling robots to reduce the reaction volume and the cost per reaction.

Our current system has several limitations, notably in the case of target proteins with a high aggregation propensity. Indeed, these proteins may be hard to express in a soluble form in CFS, making the screening process more challenging. We showed that alternative strategies, which do not overly complexify the system, could be employed to address this issue. For instance, instead of expressing these proteins in CFS, adding purified biotinylated proteins complexed with streptavidin to CF2H reactions improved the sensitivity of the assay. Future work could explore the use of optimized lysates from strains lacking reductase genes, supplemented with chaperones and isomerase that routinely express proteins once considered intractable for lysate-based CFS^73–76^.

We also found that recurrent synthesis errors in linear DNA fragments can result in false negatives (**Supplementary Note 1**). Like any method dependent on non-sequenced DNA, CF2H is highly reliant on the quality of the synthetic DNA used. Nevertheless, the generally high quality, combined with the speed and convenience offered by linear DNA, outweighs these concerns, particularly in the context of screening campaigns where libraries often contain multiple positive hits.

In terms of sensitivity and detection limits, CF2H is capable of detecting binding affinities within the nanomolar range. Regarding the upper range detection limit, our results suggest that interactions with dissociation constants above one hundred nanomolar are not reliably detectable (**Fig2f**). However, the fact that assay saturation appears to occur at very low K_d_ values, with a dynamic window between the nanomolar and picomolar ranges, suggests that CF2H is well-suited for identifying ultra-high affinity binders during screening (see **supplementary table 1** for K_d_ list). In addition to binding affinity, differences in expression levels, solubility, and stability among binders likely influence the signal output. The fact that CF2H captures all these biochemical parameters is a valuable feature, as it helps prioritize binders with favourable biophysical properties in addition to high affinity. This was successfully applied to discover a novel anti-mPD-L1 sub-nanomolar binder with immune checkpoint neutralization activity (DBP035).

Going beyond protein binder screening, we also repurposed CF2H to detect modulations of PPIs through addition of small molecules or relevant proteins. This modality could be particularly relevant for screening drugs such as competitive inhibitors or molecular glues. We also exploited this feature to repurpose CF2H as a biosensing platform and could apply it to detect relevant biomarkers such as the SARS-CoV-2 spike protein. Although CFS-based biosensors hold great promise for point-of-care diagnostics by providing affordable, field-deployable and sensitive alternatives to existing analytical methods, they remain limited by a narrow scope of detectable ligands^77^. We believe this workflow has the potential to accelerate the discovery of biosensors with new binding capabilities, either engineered from natural systems or designed entirely *de novo*.

In conclusion, we have developed a rapid and modular platform for the reliable screening of protein-protein interactions, addressing a key bottleneck in the identification of *de novo* binders. The translational potential of our platform is supported by its broad applicability—from testing small molecule modulators of PPIs and chemically inducible dimerization (CID) systems to the engineering of biosensors detecting clinically relevant targets. By enabling the development of tailored diagnostic and therapeutic tools, we envision that CF2H will offer a robust and generalizable approach for advancing applications in biomedicine and beyond.

## Methods

### Linear DNA preparation

Linear dsDNA fragments were ordered as e-blocks (Integrated DNA Technologies) or as gene fragments (Twist Bioscience) and resuspended at 10 ng/ul or 0.5 ng/μl in nuclease-free (NF) water, respectively. In the case of CF2H screening, fragments were directly PCR amplified using Platinum™ SuperFi II DNA Polymerase (ThermoFisher Scientific 12361010) following supplier’s instructions using primers described in **Supplementary Table 2.** For the rest of the experiments, PCRs were carried out on sequence-verified plasmids. Purity of PCR products were assessed by agarose gel electrophoresis (0.8% agarose, 1X TAE) and concentrations were measured using Qubit™ fluorometer dsDNA Broad Range kit (ThermoFisher Scientific Q32850). Amplified DNA fragments were normalized to 50 nM before being used in cell-free TX-TL reactions.

### Molecular cloning

Linear fragments were cloned into a low-copy pSB4k5 vector linearized by PCR using primers listed in **Supplementary Table 1** using Gibson assembly. In vitro assembled plasmids were purified using Monarch® Spin PCR & DNA Cleanup Kit (New England Biolabs,T1130S), eluted in 8 µl of NF water and 2 µl were and electroporated into homemade electrocompetent NEB10β cells (New England Biolabs). Cells were recovered in LB medium for 1 hour at 37°C 190 rpm, and subsequently plated on LB-agar supplemented with 50 μl/ml kanamycin. Finally, constructions were verified by Sanger sequencing (Eurofins Genomics).

### Cell-free extract preparation

Cell-free TX-TL extracts were prepared as previously described^18,23^. Briefly, BL21 Rosetta 2 ΔRecBCD cells grown to saturation overnight at 37°C in 2XYT-P medium were diluted 1:100 into 2L baffled flasks containing 660 mL of 2XYT-P and subsequently grown at 37°C 190 rpm until OD_600_ of 1.5-2.0. Cells were then pelleted by centrifugation, and washed twice with 200 mL of S30A buffer, before being resuspended in 0.9-1 mL of S30A per gram of dried pellet. Cells were lysed using an Avestin EmulsiFlex-C3 homogenizer at 15000-20000 psi, and the supernatant was dialysed overnight in S30B. A final centrifugation step was carried out at 12000 x g for 30 min, the supernatant was filtered through 0.22 μm, aliquoted, flash frozen, and kept at -80°C until use.

### Cell-free buffer preparation and calibration

Cell-free buffer was prepared and optimized as previously described^18,23^. For optimization of Mg- and K-glutamate concentrations, *in vitro* cell-free reactions were carried out using linear DNA driving the expression of sfGFP. First K-glutamate was fixed at 80 mM and increasing concentrations of Mg-glutamate were tested. The optimal concentration of Mg-glutamate was then used to find the optimum for K-glutamate.

### *In vitro* cell-free TX-TL reactions preparation

All cell-free reactions were prepared on ice in a final volume of 22 μl. Typically, a reaction consisted of ⅓ of bacterial extract, 5/12 of cell-free buffer, and the final ¼ of the reaction was used for DNA or drugs depending on the application. Reporter plasmid DNA was diluted to a final concentration of 5 nM from a 20X stock solution, and 2.5 nM - 5 nM of linear DNA fragments were used from 10X stock solutions or 20X stock solutions when small-molecules or proteins were spiked in. For assays involving purified proteins and small molecules, the addition was done before the reactions were started at 37°C. Then, 5 μl of each homogenized reaction were added into a low-binding 384-well plate, black walls transparent bottom and sealed with transparent adhesive film (ThermoFisher Scientific AB0558).

### Small molecule modulator assays

Caffeine (> 99.0 %, Sigma-Aldrich C0750) was resuspended in H2O at 100 mM, NVP-CGM097 (98.06%, MedChemExpress HY-15954) was resuspended in DMSO at 10mM, Venetoclax (99.95%, MedChemExpress HY-15531) was resuspended in DMSO at 10 mM, and Progesterone (99%, Sigma-Aldrich P8783) was resuspended in ethanol at 100 mM. For all assays involving small molecules, stock solutions of drugs were prepared, aliquoted, and stored at -20°C until used.

### CF2H assays with addition of purified proteins

For assays involving the addition of proteins, CF2H reactions were prepared as described above. Streptavidin (STN-N5116, Acrobiosystems), biotinylated mPD-L1 (PD1-M82F5, Acrobiosystems), biotinylated mCD47 (CD7-M82F5, Acrobiosystems), biotinylated mCTLA4 (CT4-M82F3, Acrobiosystems) and SARS-CoV2 S protein trimer (Z03501, Genscript) were resuspended according to manufacturer’s instructions, aliquoted and stored at -80°C until used. Streptavidin, mPD-L1, mCD47 and mCTLA-4 were added at concentrations of 150 nM. Complex formation with streptavidin was done by mixing proteins in a 1:1 molar ratio.

### Fluorescence Measurements

sfGFP production at 37°C was monitored using Biotek Cytation 3 or Biotek Synergy H1M plate-reader (Ex 485 nm, Em 528). To guarantee comparable measurements independently of the machine used, fluorescence intensity values were converted to FITC mean equivalent fluorescence values (MEF) as described by Jung and colleagues ^78^.

### Protein expression and purification

*De novo* protein binders DNA sequences comprising a N-terminal hexahistidine tag and C-terminal HiBit tag were cloned in a pET28a(+) vector by Genscript and transformed into homemade chemically competent *E. coli* BL21 (DE3) cells. A single colony was grown overnight in LB medium supplemented with 0.36% D-glucose 37°C, 190 rpm. Saturated overnight culture was diluted 1/50 in 500 ml of ZYM 50-52 auto induction medium and grown for 6 hours at 37°C followed by 25°C overnight at 190 rpm. Cells were pelleted by centrifugation at 6000 rpm for 25 min at 4°C. Pellets were resuspended in buffer A (50 mM Tris, 150 mM NaCl, 2 mM DTT) supplemented with lysozyme 10 μg/ml and cOmplete™ EDTA-free protease inhibitor at pH8.5 for DBP001, DBP017 and DBP035 and pH7.5 for DBP023 and stored at -80°C until further use. Cells were lysed by pulses of sonication on ice 2 x 5 min and debris were cleared by centrifugation at 18 000 rpm for 30 min at 4°C. Supernatant was loaded onto a Histrap column and eluted with a gradient of buffer B (50 mM Tris, 150 mM NaCl, 2mM DTT, 300 mM imidazole) at pH8.5 or 7.5. Fractions corresponding to the proteins of interest were pooled and dialyzed overnight in one litre of buffer A. A final size exclusion step was performed by injecting the dialyzed samples into a HiLoad ® S75 16/60 superdex column equilibrated with buffer A. Proteins were concentrated to at least 1 mg/ml if necessary and stored at -80°C.

Single domain antibody VHH-1D with N-terminal PelB periplasmic secretion signal a C-terminal hexahistidine tag followed by a HiBit tag was cloned in a pET22b vector by Genscript and transformed into homemade chemically competent *E. coli* BL21 (DE3). Expression was performed following the same procedure as for the de novo binders. Purification of periplasmic VHH was performed as previously described ^79^. VHH-1D was eluted in PBS pH7.4, concentrated to 1 mg/ml and stored at -80°C.

### *In silico* binder design with Bindcraft

Binders for mouse PD-L1 (mPD-L1) were designed using BindCraft ^10^ V1.2.0, available at <u>GitHub</u>. The software was executed on an NVIDIA A6000 GPU, using the mouse PD-L1 structure (PDB: 6SRU) as input and targeting residues 54, 56, 58, 60, 113, and 115, with binder size defined ranging from 65 to 155 residues. BindCraft was configured with the following settings: `settings_filters: default_filters` and `setting_advanced: default_4stage_multimer`. A total of 100 designs were generated, ranked by decreasing interface predicted template modelling score (i_pTM), and the top 36 designs were selected for experimental screening.

### HiBit-Blot

Samples from CF2H reactions ran for 4 hours at 37°C were mixed in 2X loading buffer (125 mM Tris pH6.8, 20% glycerol, 4% SDS, 250 mM DTT) and loaded onto a NuPAGE™ 4-12% bis-tris in 1X MES buffer and ran at 220V for 40 min. Blotting and revelation were performed as recommended by Promega. Proteins were transferred onto a 0.2 μm nitrocellulose membrane using a Biorad Trans-Blot Turbo Transfer System on the Turbo settings. The membrane was first washed in TBS-T (20 mM Tris, 150 mM NaCl, 0.1% Tween pH7.4) before being submerged in blotting buffer solution (1X) supplemented with the LgBit protein for 1 hour at RT. Finally, Nano-Glo® luciferase assay substrate was diluted 500 times directly into the blotting solution and luminescence was imaged immediately using Amersham™ 600 imager.

### HiBit-ELISA

White opaque 96-well microplates (Thermo Fisher Scientific) were coated overnight at 4C using 100 μl of mPD-L1 (PD1-M5251, Acrobiosystems) or Bovine Serum Albumin (BSA) (10735078001, Merck) solutions at 1 μg/ml and 2 μg/ml per well, respectively. Wells were then washed three times in PBS-T (137 mM NaCl, 2.7 mM KCl, 10 mM Na2HPO4, Tween-20 0.05%, pH 7.4), before a blocking step in 5% non-fat milk in PBS-T for 1 hour at RT. Blocking solution was discarded and wells washed 4 times with PBS-T. Serial dilutions of purified mPD-L1 binders tagged with a HiBit were then added to the wells and incubated for 1 hour at RT. Finally, binders were discarded and wells washed 8 times with PBS-T, the HiBiT reagent mix (Nano-Glo ® extracellular buffer, LgBiT, Nano-Glo® luciferase assay substrate) was diluted 1:1 in PBS-T before adding 100 μl in each well. The luminescent signal was measured using a Biotek Cytation 3 plate-reader.

### Surface Plasmon Resonance

SPR experiments were carried out using a Biacore T200 instrument (Cytiva) at 25°C with a PBS buffer containing 0.05% (v/v) Polysorbate 20 as the running buffer. Anti-human Fc antibodies (Cytiva) were immobilized on a CM5S sensor chip via amine coupling according to the manufacturer’s instructions. The mPD-L1-Fc protein (PD1-M5251, Acrobiosystems) was captured on the chip surface to a level of 250–400 RU by injecting at 10 µL/min for 180 seconds. Kinetic binding assays of de novo binders (DBP001, DBP023, and DBP035) were performed using multi-cycle kinetics. A two-fold serial dilution series of the mini-protein binders (0.625 – 20 nM) was injected over the captured PD-L1-Fc surface at 100 µL/min for 180 seconds, followed by a 400-second dissociation phase with a running buffer. The sensor surface was regenerated after each cycle by injecting 3 M MgCl₂ at 30 µL/min for 30 seconds. Data were processed using Biacore T200 Evaluation Software version 3.2. Double referencing was applied before fitting the binding curves to a 1:1 Langmuir binding model to determine kinetic parameters.

### PD1/PD-L1 immune checkpoint blockade assay

Neutralizing activity of anti-mPD-L1 binders was assessed using a mouse PD-1/PD-L1 Blockade Bioassay (Promega, CS303201) according to the manufacturer’s protocol. Briefly, PD-L1 aAPC/CHO-K1 cells were plated into a 96-well plate and incubated for 16 h. Binders and anti-mPD-L1 monoclonal antibody solutions (BioLegend, 124301) solutions were prepared in PBS by doing 2.5-fold serial dilutions (20 μM to 134 pM for binders and 32 nM to 134 pM for mAb). After 6 h of incubation, the Bio-Glo™ reagent was added and luminescence was measured after 15 min using a BioTek Cytation 3 plate reader.

## Acknowledgements

We thank members of our groups and institutes for fruitful discussions and feedback. We are grateful to P. Voyvodic, H.J. Chang, P. Soudier, and A. Levrier for their contributions to the initial iteration of the split-DBD and linear CFS systems in our lab. We thank JL Pons and F Wollscheidt for help with our computing infrastructure. JB thanks INSERM and the Bettencourt-Schueller Foundation for their continuous support. This work was supported by an INSERM grant for the development of a Technological Research Accelerator in Synthetic Biology (ART-synbio). JC salary was funded by the European FET-Open grant BioCellPhe to JB (Grant number: 965018). The CBS acknowledges support from the French Infrastructure for Integrated Structural Biology (FRISBI; ANR-10-INSB-05-01). SPR experiments were carried out using the facilities of the Montpellier Proteomics Platform (PPM, BioCampus Montpellier), a member of the Proteomics French Infrastructure (ProFI).

## Authors contributions

JB and JC conceived the study. JC, PM, ADV, AB, JE and G.N. conducted experiments and analyzed data. AC and RJR contributed data for MdM2 binders and insight into structural and biophysical analyzes. MCG supervised protein production and provided expertise in structural biology and biophysics. JB and JC wrote the manuscript with input from all authors. All authors read and approved the final version of the manuscript.

